# Amphipathic Fluorescent Dyes for Sensitive and Long-Term Monitoring of Plasma Membranes

**DOI:** 10.1101/2020.11.16.379206

**Authors:** Masataka Takahashi, Ryo Seino, Takatoshi Ezoe, Munetaka Ishiyama, Yuichiro Ueno

## Abstract

The plasma membrane (PM) plays a critical role in many cellular processes, and PM dysfunction is a key biomarker related to the cell status and several diseases. Imaging techniques using small fluorescent probes have become increasingly important tools for visualizing living cells, particularly their PMs. Among the commercially available PM-specific probes, PKH dyes are widely used; however, the utility of these dyes is limited by their short membrane retention times and high cytotoxicity. Herein, PlasMem Bright Green and Red are implemented as new PM-specific fluorescent probes, which employ a polycyclic aromatic fluorophore to improve their retention ability and a strong acid moiety to reduce their transmembrane diffusion and cytotoxicity. We demonstrate that the long retention and low cytotoxicity of the PlasMem Bright dyes enable them to be applied for observing neuronal PMs and monitoring PM dynamics involving the endocytic pathway. Furthermore, we successfully detected mitochondria in nerve axons over long periods using PlasMem Bright dyes. Finally, the combined use of exosome staining probes and PlasMem Bright dyes allowed clear visualization of the endocytic pathway.

**TOC graphic:** 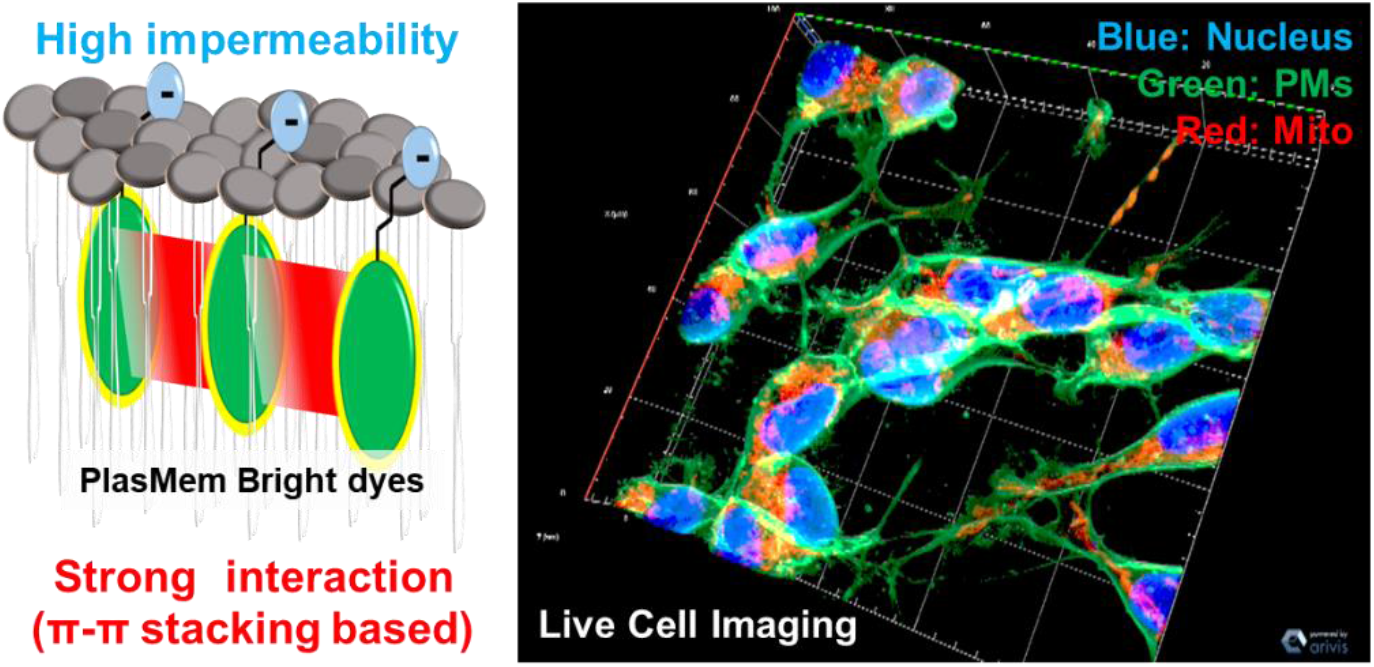

## INTRODUCTION

The plasma membrane (PM) consists of a lipid bilayer, which separates the intracellular environment from extracellular spaces. Consequently, the PM plays an essential role in many cellular phenomena, such as cell migration,^1,2^ stretching,^3^ and signaling cascades.^4^ In addition, because of the implication of PM dysfunction in so many diseases, it is crucial to establish a fundamental understanding of PM morphology and dynamics; however, comprehensive investigations aimed at elucidating these aspects have not yet been conducted.^5,6^ Generally, methods for monitoring PM morphology and dynamics are either based on labeling the PM with small fluorescent molecules, expressing fluorescent proteins targeted to the PM via plasmid transfection,^7^ or immunostaining the PM with antibodies.^8^ The transfection and immunostaining techniques face major limitations, such as requiring the stable expression of the fluorescent protein and having limited applicability for fixed cells. In contrast, small fluorescent molecules are widely used because they can be added to live cells,^9^ and the most commonly-used small fluorescent molecules are PKH dyes (named for the scientist who developed them, Paul Karl Horan).^10^ Although PKH dyes contain a cationic moiety and C-18 hydrocarbon chains, they have not been designed or functionalized specifically for PM monitoring. Additionally, because of the high hydrophobicity of these dyes, a washing step is necessary before the stained cells can be observed under a microscope.^11^ These characteristics of PKH dyes limit their applicability for observing PM dynamics, including the endocytosis pathway,^12^ in high contrast. Therefore, there is a demand for developing a new PM-specific probe, which is a great technical challenge but also an essential and urgent necessity.

Herein, we report the application of newly-synthesized fluorescent probes, called PlasMem Bright, which employ (i) polycyclic aromatic fluorophores to improve the retention ability of the probes when interacting with the PM and (ii) an anionic moiety to reduce their cytotoxicity. This work indicated that the PlasMem Bright dyes are more water-soluble and can be retained in the PM for longer in comparison with PKH dyes. Moreover, for nerve axonal imaging, the PlasMem Bright dyes allowed us to capture high-contrast images of PMs, with very low toxicity to the cells compared with highly-cytotoxic CellMask dyes. Furthermore, we successfully detected mitochondria in nerve axons using PlasMem Bright dyes and mitochondrial staining dyes. These results encouraged us to further investigate the PlasMem Bright dyes for other applications, including monitoring the endocytic pathway using an exosome staining probe. Indeed, the combined use of an exosome staining probe and PlasMem Bright dyes allowed for clear observation of the endocytic pathway.

## Materials and Methods

### Reagents

The PKH26 Fluorescent Cell Linker Mini Kit was purchased from Merck-Sigma Aldrich. The CellMask Green and Orange were purchased from Thermo Fisher Scientific. MitoBright LT Green, Red and ExoSparkler Exosome Membrane Labeling Kit-Red were obtained from Dojindo Laboratories.

### Cell culture

HeLa, SH-SY5Y, and WI-38 cells were cultured in Minimum Essential Media (MEM, Thermo Fisher Scientific) supplemented with 10% (v/v) fetal bovine serum, 1% non-essential amino acid, 1% penicillin, and 1% streptomycin. All cells were maintained in a humidified 5% CO_2_ incubator at 37 °C. Jurkat cells were cultured in Roswell Park Memorial Institute (RPMI, Thermo Fisher Scientific) supplemented with 10% (v/v) fetal bovine serum and 1% penicillin.

### Confocal Imaging

Cells (HeLa, SH-SY5Y, WI-38, Jurkat; 2.5 x 10^5^) were seeded on the ibidi-8well plate for 24 h before imaging. After removal of the medium, the cells were stained in culture medium containing 0.5% DMSO and dye in a CO_2_ incubator under the following conditions: 5 μmol/l PlasMem Bright dyes for 30 minutes; 5 μmol/l PKH26 for 30 minutes; 100-fold dilution of CellMask dyes for 30 minutes; 0.1 μmol/l MitoBright LT dyes for 30 minutes. The cells were then washed with Hanks’ Balanced Salt Solution (HBSS) to remove the free dye and kept in culture medium for imaging. The imaging data were obtained with LSM 800 confocal laser scanning microscopy (ZEISS): excitation at 488 nm and an emission filter of 500-550 nm for PlasMem Bright Green, CellMask Green, and MitoBright LT Green: excitation at 561 nm and an emission filter of 560-650 nm for PKH26, CellMask Orange, and labeled-exosome: excitation at 561 nm and an emission filter of 560-700 nm for PlasMem Bright Red.

## RESULTS

### Investigation of the PM staining mechanism using PlasMem Bright dyes

PlasMem Bright Green and Red are amphipathic dyes, which contain a polycyclic aromatic fluorophore and a strong acid moiety, and function as green- and red-emitting fluorescent probes, respectively. The fluorescence signal of PlasMem Bright dyes is turned ON in nonpolar environments, such as a lipid bilayer membrane, and turned OFF in aqueous media, such as extracellular environments. Introducing an electron-donating piperazine moiety into the PlasMem Bright Red fluorophore produced a donor-π-acceptor-type fluorophore that exhibits environment-sensitive emission due to twisted intramolecular charge transfer (ICT) in the excited state.^13,14^ This characteristic led PlasMem Bright Red to provoke a bathochromic shift in both the absorbance and emission spectra, which solidifies its use in the red channel, with limited crosstalk into the green channel. These features allow the PM morphology to be observed in high contrast.

These dyes exhibit affinities for PMs because of two synergic interactions. Specifically, the enhanced hydrophobicity of the polycyclic aromatic fluorophore component promotes interactions with the hydrophobic tails of the lipids, and the π-π stacking between fluorophore moieties contributes to the longer retention of the dyes in the PM by inducing strong intermolecular interactions. Additionally, the strong acid component repels the polar heads of the PM (Figure 1).

**Figure 1.**
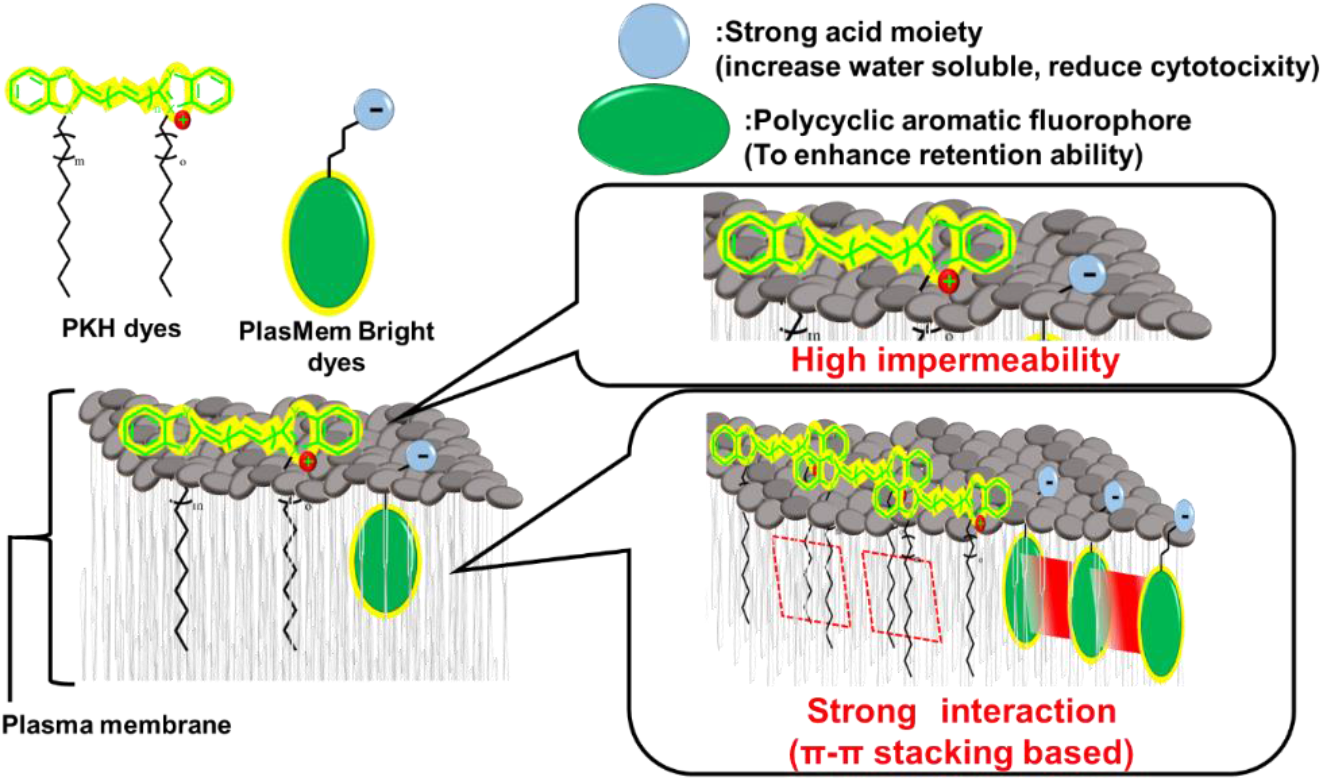
Mechanism of PM staining.

Compounds A and B were synthesized as references for PlasMem Bright dyes and were evaluated using microscopic analysis. In these molecules, the strong acid moiety was replaced with a pentyl group to increase the hydrophobicity. Theoretically, the lack of strong acid moiety in the chemical structures of Compounds A and B should cause them to diffuse more easily through the PM relative to the PlasMem Bright dyes (Figure 2).

**Figure 2.**
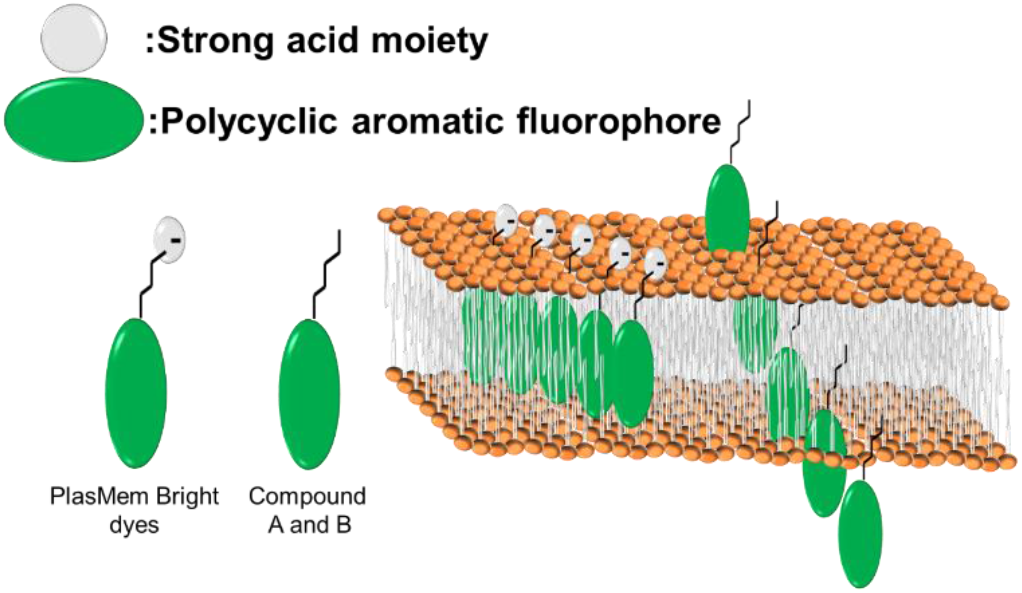
Structure of PlasMem Bright dyes and Compounds A and B (left), and these compounds’ proposed interactions with the PM (right).

The fluorescence signals of PlasMem Bright dyes were exclusively observed at the PM, whereas the fluorescence signals of Compounds A and B were observed at the PM and with intracellular components, indicating that the strong acid moiety on PlasMem Bright dyes is critical for PM staining (Figure 3).

**Figure 3.**
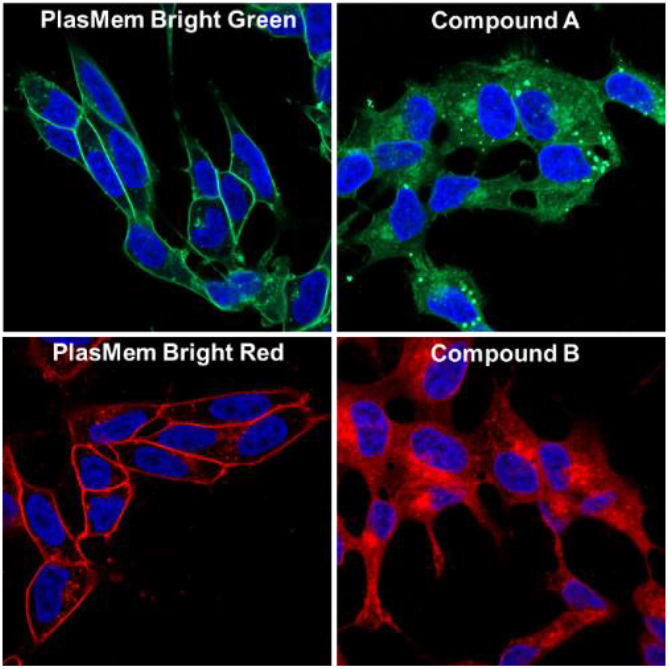
Fluorescence images of HeLa cells stained with PlasMem Bright dyes (left) and Compounds A and B (right).

To further investigate the specificity of PlasMem Bright dyes for the PM, co-localization experiments using either the PKH26 dye (Merck-Sigma Aldrich) or CellLight Plasma Membrane-RFP (Thermo Fisher Scientific) were performed. As shown in Figure 4, the PlasMem Bright dyes demonstrated good co-localization with both the PKH26 dye and the CellLight Plasma membrane-RFP, indicating high PM selectivity.

**Figure 4.**
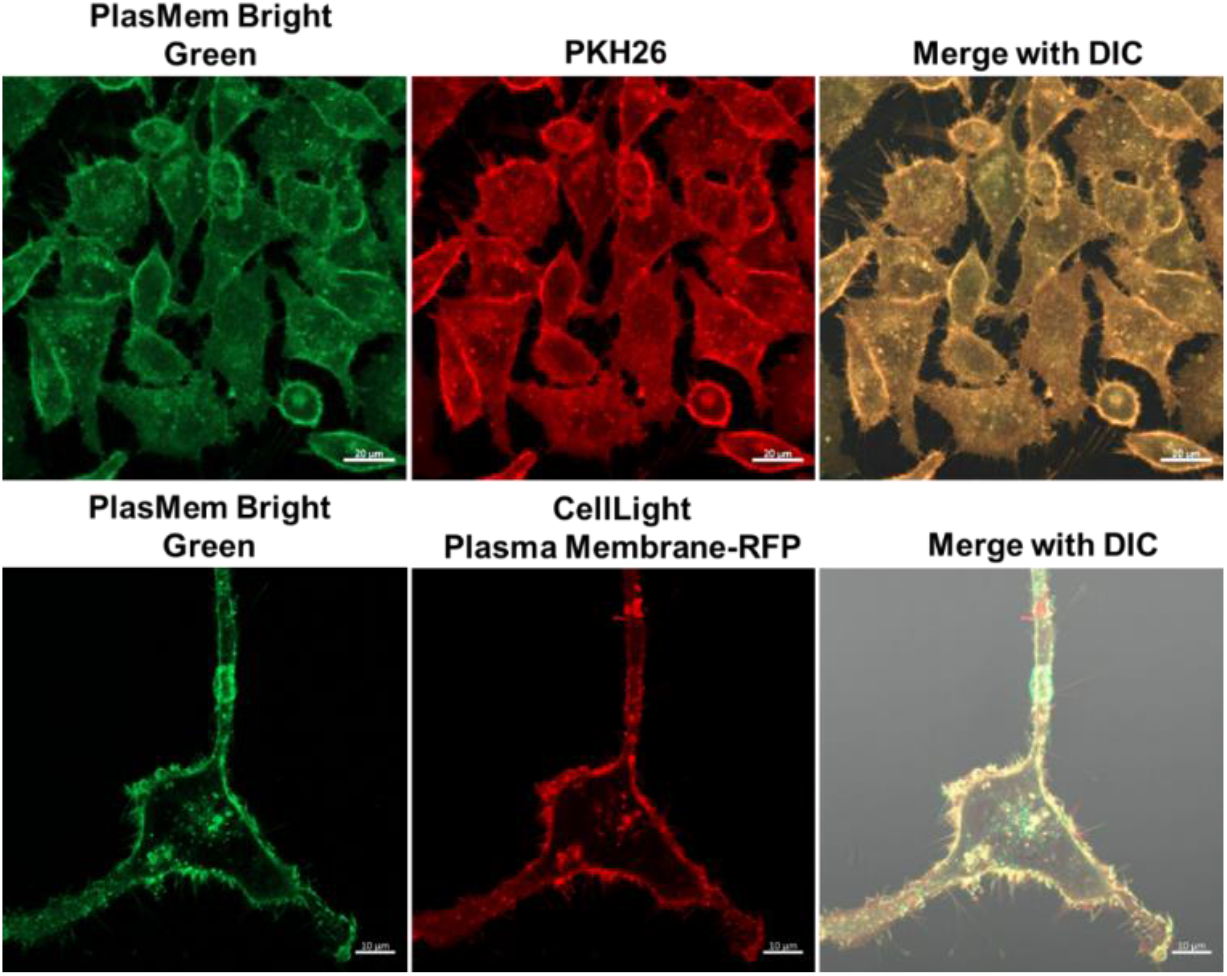
Fluorescence images of HeLa cells stained with PlasMem Bright Green (left), PKH26 (center of upper column), and CellLight Plasma Membrane-RFP (center of lower column).

The fluorescent microscopic 3D analysis was carried out using PlasMem Bright Green-stained HeLa cells to confirm the PM-specific staining ability of PlasMem Bright dyes (Figure 5).

**Figure 5.**
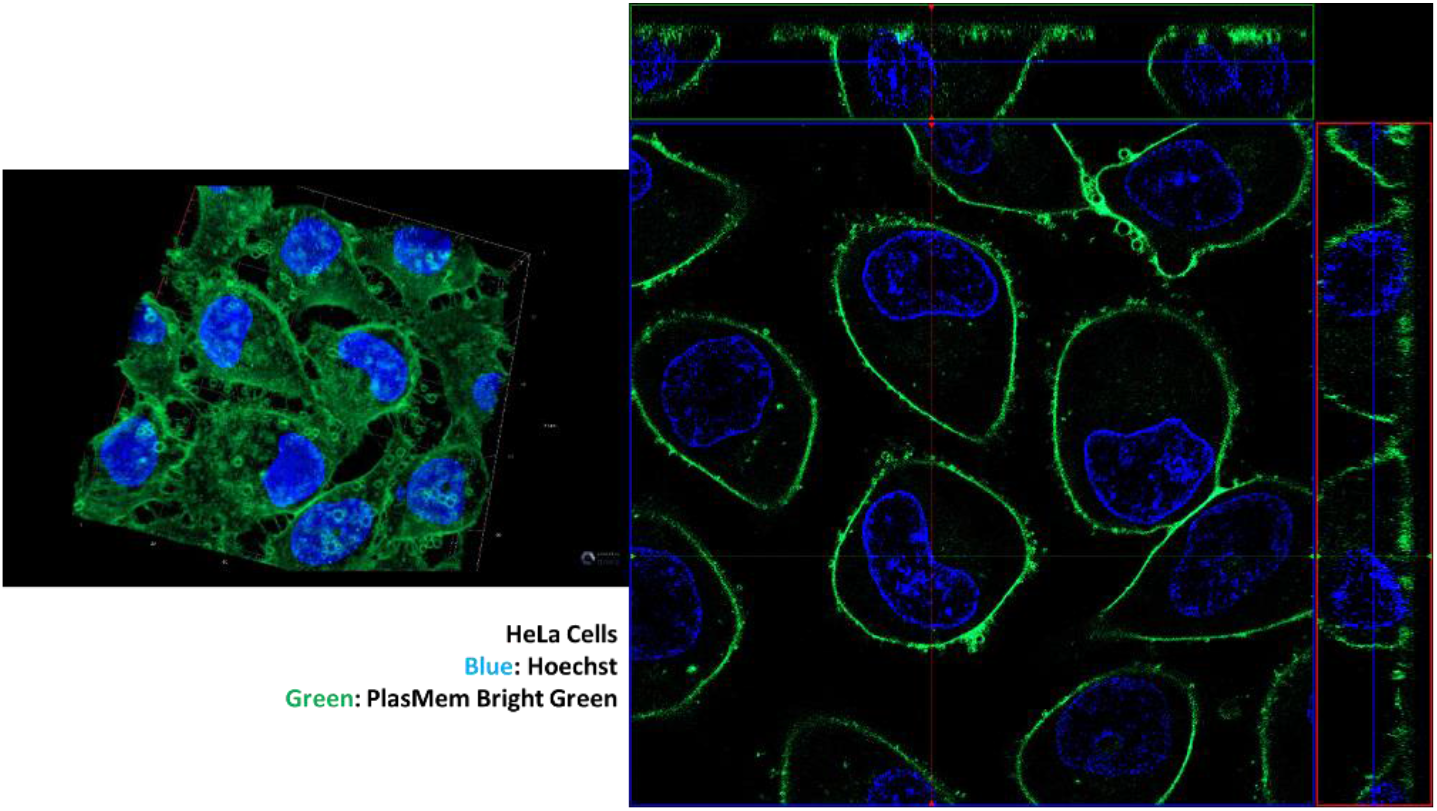
3D image (left) and 3D ortho data (right) of HeLa cells stained with PlasMem Bright Green and Hoechst 33342.

PM staining using the PlasMem Bright Green dye without any washing steps was also carried out using SH-SY5Y, Jurkat, and WI-38 cells, with high specificity.

**Figure 6.**
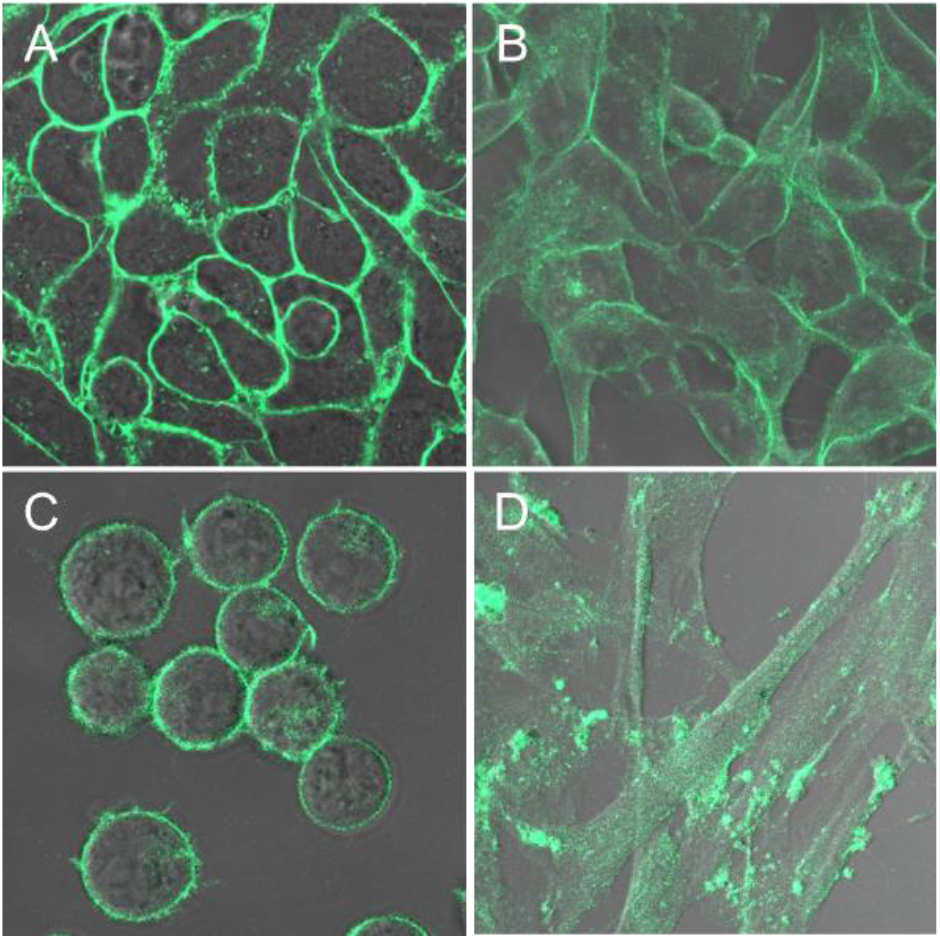
Images of live cells from four different cell lines stained with 5 μM of PlasMem Bright Green. (A) HeLa cells, (B) SH-SY5Y cells, (C) Jurkat cells, (D) WI-38 cells.

The retention of PlasMem Bright Green in the PM was examined in comparison with PKH26. HeLa cells were co-stained with PlasMem Bright Green and PKH26 and then washed with culture medium. After 24 and 48 hours of incubation, the stained HeLa cells were observed under a confocal microscope. As shown in Figure 7, PlasMem Bright Green exhibited greater retention in the PM compared with PKH26, even after 48 hours of incubation.

**Figure 7.**
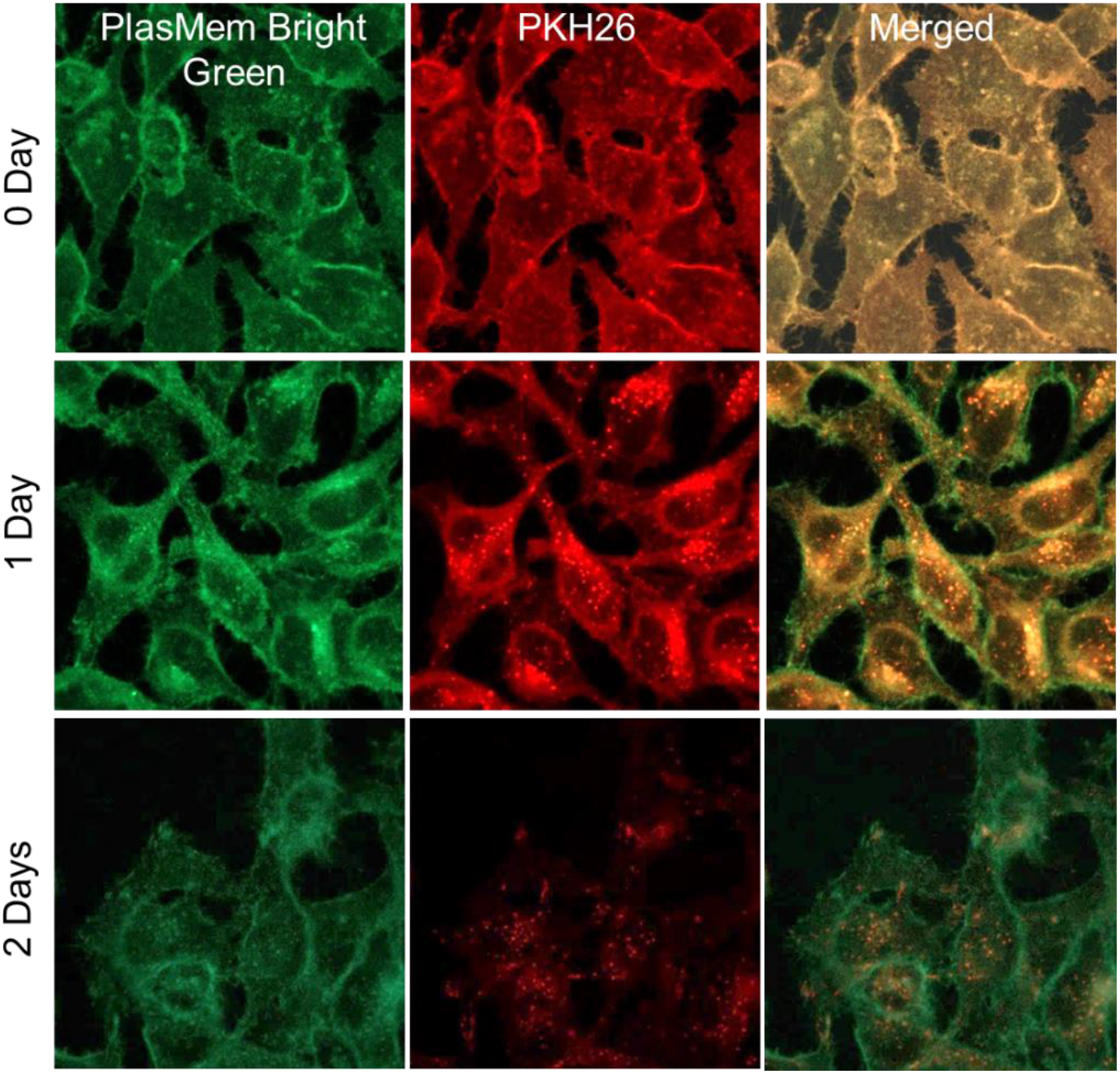
Comparison of staining performance in HeLa cells over 48 hours: PlasMem Bright Green (left) and PKH26 (center).

### Monitoring mitochondria in nerve axons

To explore additional applications for PlasMem Bright dyes, these compounds were tested towards monitoring mitochondria in nerve axons. To our knowledge, there is currently no effective method for monitoring mitochondria in living nerve axons, even though mitochondrial dysfunction in nerve axons has been widely studied as a promising target for understanding neurodegenerative diseases.^15^

The neurons were differentiated from SH-SY5Y cells in DMEM/F11 medium (10 μmol/L retinoic acid, 1% fetal bovine serum), and the PlasMem Bright dyes were used at a concentration of 5 μmol/L. To stain the mitochondria, MitoBright LT dyes (Dojindo Laboratories) were used at a concentration of 0.1 μmol/L, and Hoechst 33342 was used at a concentration of 5 μg/mL. As references, CellMask Green and Orange (Thermo Fisher Scientific) were used at a concentration of 100-fold dilution with a culture medium. After staining, the cells were observed under a confocal microscope.

Nerve axons stained with CellMask Orange shrank in size after the staining, likely because of the high cytotoxicity of this dye (yellow arrows in image B of Figure 8). However, mitochondria in the tubular structures of the nerve axons were clearly detected by PlasMem Bright dye and MitoBright LT dye staining (Figure 8).

**Figure 8.**
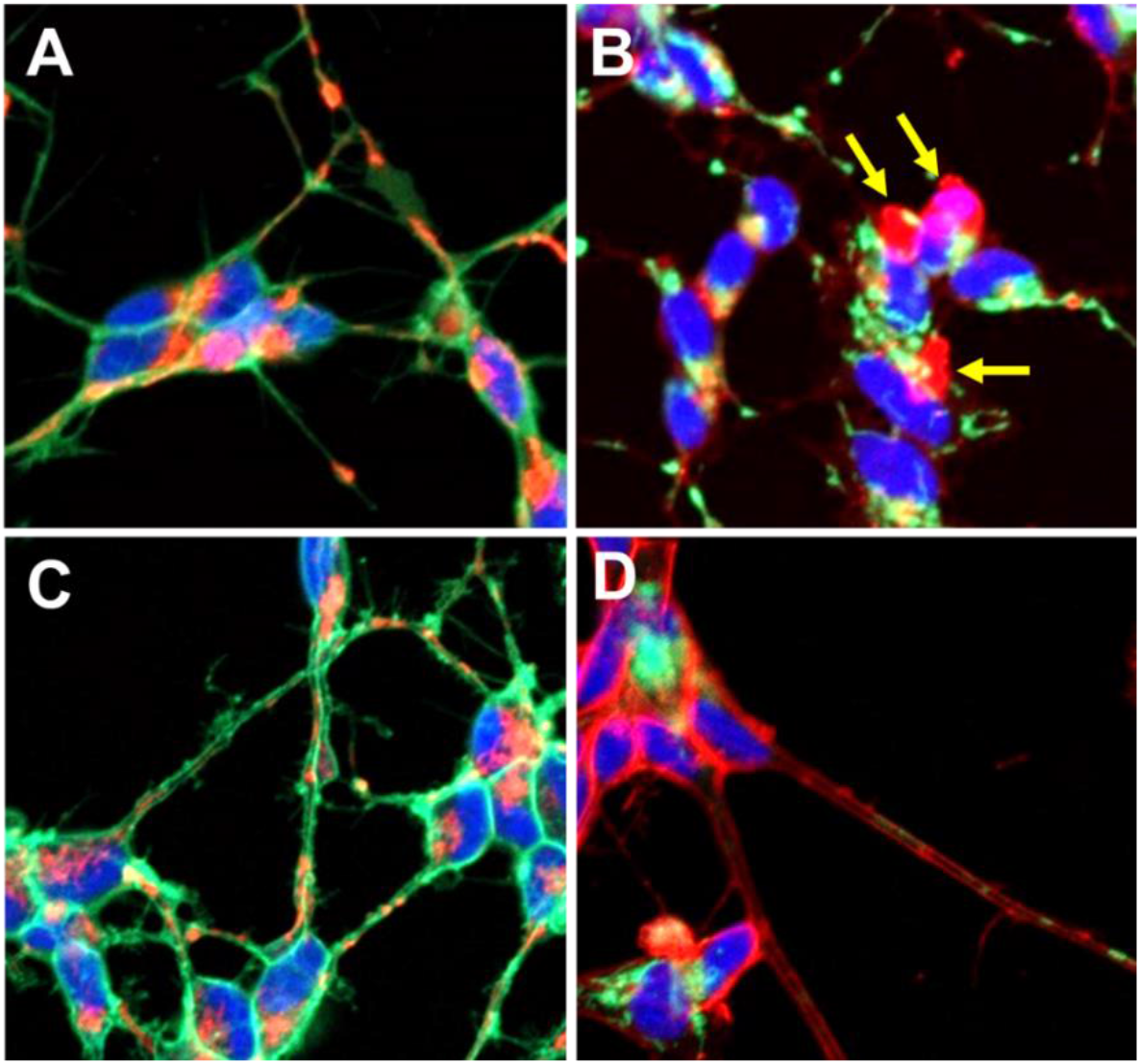
Multi-colored images of plasma membranes and mitochondria in the differentiated SHSY-5Y cells stained with PM dyes, MitoBright LT dyes, and Hoechst 33342. (A) Blue: Hoechst 33342, Green: CellMask Green, Red: MitoBright LT Red; (B) Blue: Hoechst 33342, Green: MitoBright LT Green, Red: CellMask Orange; (C) Blue: Hoechst 33342, Green: PlasMem Bright Green, Red: MitoBright LT Red; (D) Blue: Hoechst 33342, Green: MitoBright LT Green, Red: PlasMem Bright Red.

### Monitoring exosome uptake through the endocytic pathway

The PM dynamics of HeLa cells were investigated using PlasMem Bright Green and ExoSparkler dye (Dojindo Laboratories)-labeled exosomes. Exosomes are incorporated into the cells via endocytosis,^12^ so using dyes to individually stain the exosome and the membrane two different colors should allow clear observation of the incorporated exosome (stained red with ExoSparkler dye) at the site of endocytosis in the membrane (stained green with PlasMemBright dye) in the colored fluorescence image. To confirm this, ExoSparkler dye-labeled exosomes were added to PlasMem Green dye-stained HeLa cells. Following overnight incubation in a controlled environment (37 °C, 5% CO_2_), the cells were observed using a confocal microscope. As shown in Figure 9, the exosomes taken up via endocytosis were successfully detected using PlasMem Bright Green and ExoSparkler Red.

**Figure 9.**
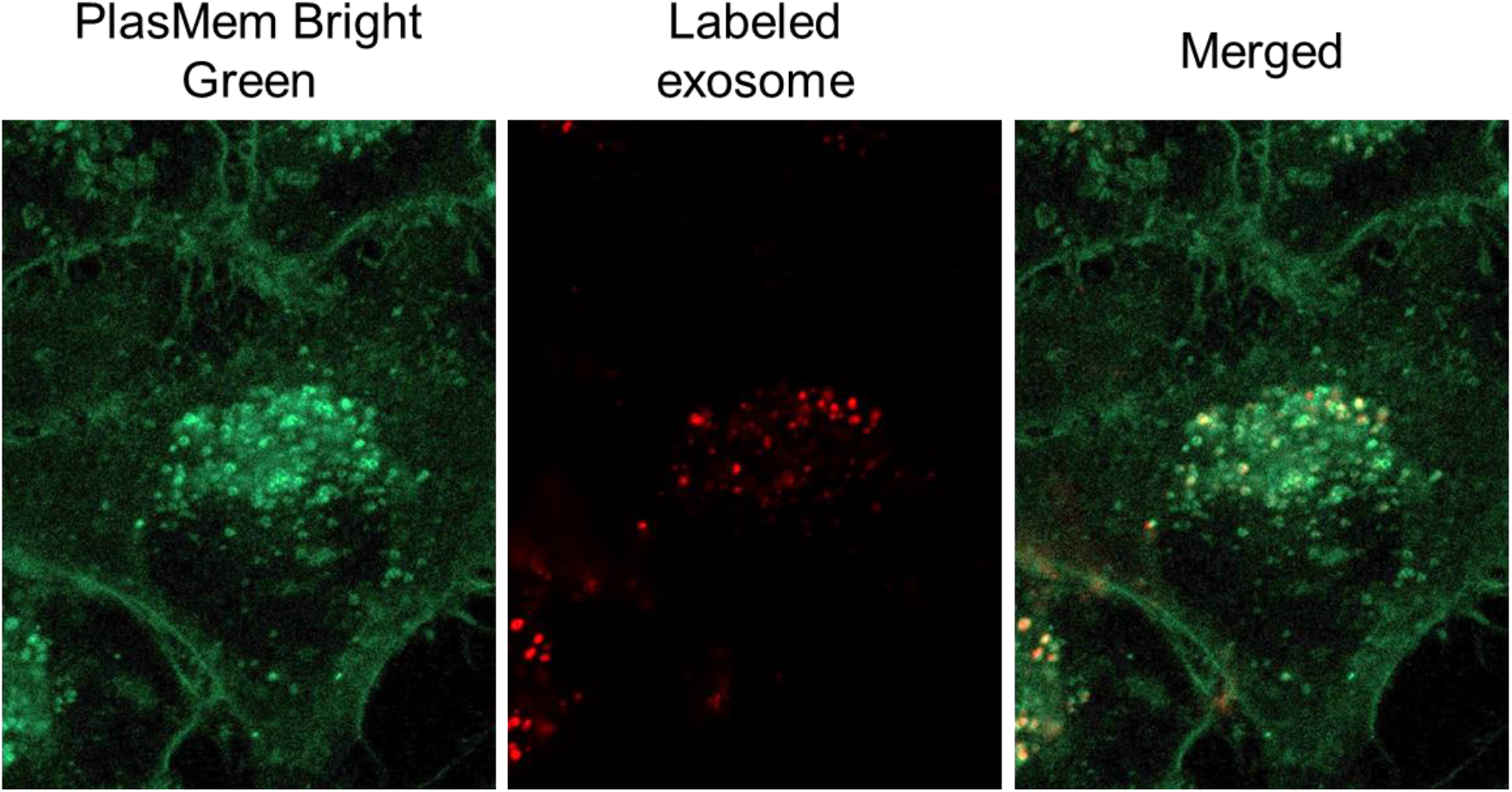
Images of fluorescent-labeled exosome in integrated plasma membrane. Green: PlasMem Bright Green, Red: Labeled exosome using ExoSparkler Red.

## DISCUSSION

In this study, we developed and successfully implemented a molecular tool for monitoring nerve axons and the endocytic pathway in living cells. The fluorescent PlasMem Bright dye molecules were designed to overcome the three principal issues facing commercially available PM staining dyes, namely, cytotoxicity, low membrane retention, and water insolubility. According to our experimental results, PKH26 suffered from a short retention time in the PM (Figure 7), and CellMask dyes exhibited high cytotoxicity towards neurons (Figure 8). We propose that the different results between PlasMem Bright dyes and PKH dyes were due to the distinct chemical structures of each dye. Specifically, introducing the amphipathic moieties in PlasMem Bright dyes represented a preferred design strategy for PM staining in living cells (Figure 1). PlasMem Bright dyes (i.e., long retention in the PM, low cytotoxicity, and high-water solubility) allowed us to observe nerve axons and the endocytic pathway in high contrast fluorescence images. Furthermore, the combined use of PlasMem Bright dyes and ExoSparkler dyes enabled us to simultaneously monitor the PM and exosome dynamics (Figure 9).

## Summary

PlasMem Bright dyes, which are newly-developed fluorescent probes, are highly water-soluble, and they exhibited excellent membrane retention abilities and negligible cytotoxicity during our experiments. Use of these PlasMem Bright dyes allows clear visualization of PM dynamics and nerve axons, with low cytotoxicity in the long-term. Because PM dysfunction is implicated in many diseases, it is important to understand fundamental PM morphology and dynamics without inducing cytotoxicity over an extended period of time. The findings reported herein establish a basis on which to develop and advance critical PM research.

## ACKNOWLEDGMENTS

We thank Suzanne Adam, Ph.D., from Edanz Group (https://en-author-services.edanzgroup.com/ac) for editing a draft of this manuscript.

